# Genome content in the non-model ciliate *Chilodonella uncinata*: insights into nuclear architecture, gene-sized chromosomes among the total DNA in their somatic macronuclei during their development

**DOI:** 10.1101/2024.11.13.623465

**Authors:** Ragib Ahsan, Xyrus X. Maurer-Alcalá, Laura A. Katz

**Author notes:** Corresponding author Laura A. Katz.

## Abstract

Ciliates are a model lineage for studies of genome architecture given their unusual genome structures. All ciliates have both somatic macronuclei (MAC) and germline micronuclei (MIC), both of which develop from a zygotic nucleus following sex (i.e., conjugation). Nuclear developmental stages are not as well explored among non-model ciliate genera, including *Chilodonella uncinata* (Class-Phyllopharyngea), the focus of our work. Here, we characterize nuclear architecture and genome dynamics in *C. uncinata* by combining DAPI (4′,6-diamidino-2-phenylindole) staining and fluorescence *in situ* hybridization (FISH) techniques with confocal microscopy. We developed a telomere probe for staining alongside DAPI, which allows for the identification of fragmented somatic chromosomes among the total DNA in the nuclei. We quantify both total DNA and telomere-bound signals to explore changes in DNA content and chromosome maturation across *Chilodonella*’s nuclear life cycle. Specifically, we find that MAC developmental stages in the ciliate *C. uncinata* are different than the data reported from other ciliate species. These data provide insights into nuclear dynamics during nuclear development and enrich our understanding of genome evolution in non-model ciliates.

## Introduction

Ciliates are a ∼1 billion year old clade of diverse eukaryotic microorganisms (Chen et al. 2015; Howard-Till et al. 2022; Parfrey et al. 2011; Philippe et al. 2000). One of the key characteristics of ciliates is the presence of dimorphic nuclei within a single individual, where ciliates possess at least one somatic macronucleus (MAC) and at least one germline micronucleus (MIC) (Ahsan et al. 2022; Rzeszutek et al. 2020). Somatic macronuclei are highly polyploid with active gene expression throughout the life cycle (Bétermier et al. 2023; Chalker et al. 2013; Duharcourt et al. 2009; Raikov 1995). The germline micronucleus is diploid and remains quiescent (i.e. DNA is in a heterochromatic state) during asexual growth. Germline micronuclei generate gametic nuclei through meiosis, which are exchanged during conjugation (Chalker et al. 2013; Jönsson 2016; McGrath and Katz 2004; Pilling et al. 2017; Prescott 1994; Raikov 1969, 1982). The number, shape, and structure of MACs and MICs varies among species in ciliates (Ahsan et al. 2022).

Ciliates have unusual nuclear structures and complex life cycles in which they alternate between asexual division and sex through conjugation. During asexual reproduction, most ciliates reproduce by binary fission during which micronuclei divide by mitosis and polyploid macronuclei divide by amitosis (Bellec et al. 2014). During sexual reproduction, ciliates go through conjugation, where germline micronuclei produce haploid products through meiosis which are exchanged with each other (Bellec et al., 2014; Bradbury, 1966; H. Darby, 1930; Morgens et al., 2014; I. B. Raikov, 1982), and then fuse to form a zygotic nucleus. This zygotic nuclei divides mitotically, with one daughter nucleus developing into a new somatic macronuclei and the other becoming a new germline micronuclei (Ahsan et al., 2022). Though nuclear structure and genome dynamics are well explored in several model ciliates (*Tetrahymena, Oxytricha*, and *Paramecium*, Ammermann et al. 1974; Chalker 2008; Chalker et al. 2013; Ishida et al. 1999; Lipps and Eder 1996; Postberg et al. 2001; Stevenson and Lloyd 1971; Zhang et al. 2023), the extent of how well these processes reflect the bulk of ciliate diversity remains poorly understood (Russell et al. 2017; Yan et al. 2017; Maurer-Alcalá et al. 2018; Zheng et al. 2021).

Here, we focus on *Chilodonella* (class-Phyllopharyngea) as it is both cultivable and among non-model ciliates whose nuclear architecture is yet to be fully explored. *Chilodonella*’s somatic nuclear architecture is quite distinct, harboring dense DNA-rich sphere-like heteromeric structures (Maurer-Alcalá and Katz 2016) that surround a DNA-poor center. The density and characteristics of these DNA-rich ‘spheres’ varies across *Chilodonella*’s nuclear life cycle (Bellec et al. 2014; Maurer-Alcalá and Katz 2016).

There are some prior studies that focused on describing *Chilodonella*’s MAC development using light and electron microscopy (Pyne 1979, 1978; Pyne et al. 1974), its lifecycle (Bellec et al. 2014), and aspects of its genome biology and development (Maurer-Alcalá & Katz, 2016; Maurer-Alcala et al. 2018). Here, we propose a revised life cycle for the non-model freshwater ciliate *Chilodonella uncinata* based on modern fluorescence-based descriptions of the macronuclear developmental stages. By using DAPI (4′,6-diamidino-2-phenylindole) and fluorescence i*n situ* hybridization (FISH) using a probe designed to target the telomere sequences of this species, we are able to explore changes in total DNA content and somatic chromosome maturation across macronuclear developmental stages. We also evaluate variation in nuclear architecture during conjugation, and relate nuclear data to estimates of cell size. Overall, this study contributes to our understanding of nuclear life cycles, and particularly in the development of somatic macronuclei, in non-model ciliates.

## Methods

### Cell culture and maintenance

Cultures of *Chilodonella uncinata* (strain ATCC PRA-257), were originally isolated from a sample from Poland in 2014 and were maintained in Volvic water (bottled Volvic natural spring water) with autoclaved rice grains to support bacterial (i.e. prey) growth. Cultures of *C. uncinata* were maintained in 6-well plates, transferring 200 μl of cells into new wells with some Volvic water every 3-5 days. All cultures were maintained at room temperature and in the dark. We used a brightfield microscope (Olympus CKX31) to maintain the cultures and isolate cells for FISH experiments.

### Cell isolation for the experiment and fixation

Cells isolated from the 6-well plate wells were transferred to the Superfrost slides (Fisherbrand™ Superfrost™ Plus Microscope Slides; Catalog No. 22-037-246) using a 200 μl micropipette. Before placing the isolated cells on the superfrost slides, a square shaped boundary was drawn on the slides using a hydrophobic pen (Cole-Parmer Hydrophobic Barrier PAP Pen; Catalog No.NC1882459). Cells were left on the superfrost slides at room temperature for 20 minutes. Afterwards, a solution of 4% paraformaldehyde (PFA, product ID: J19943-K2, lot# 210699) in PBS (product code: 1003127976, lot# SLCH0992) was added at a final concentration of 2% and incubated at room temperature for 20 minutes. The majority of the liquid was then drawn off the slides, followed by three washes with 1x PBS.

### Fluorescence *in situ* hybridization

We performed fluorescence *in situ* hybridization (FISH) to localize the gene-sized chromosomes in the macronuclei in *Chilodonella uncinata*. We designed the oligonucleotide telomere probe using the direct telomeric repeats (C_4_AAA_3_)_3_. The probe was labeled with Alexa Fluor 488 fluorescent dye at the 5’ end (5’-CCCCAAACCCCAAACCCCAAA-3’).

Following fixation and washing, cells were permeabilized using 0.5% Triton X-100 for 20 minutes at room temperature, then washed twice with 1x PBS and once with 4x saline-sodium citrate (SSC). Cells were equilibrated in a pre-hybridization buffer (50% formamide, 2X SSC, and nuclease-free water (NFW) for 30 minutes at room temperature. The hybridization buffer was prepared by mixing 10μM telomere probe, 50% formamide, 4X SSC solution, and NFW. This hybridization mix was denatured at 98°C for 5 minutes and snap cooled on ice. 20 μl of hybridization buffer was added to the cells, which were incubated at 75°C for 5 minutes, then overnight at 37°C. The next day, slides were washed with 2x saline-sodium citrate for 15 minutes at 37°C, then 1x saline-sodium citrate (SSC) for 15 minutes at 37°C in a water bath, with a final wash with 1x SSC at room temperature on the bench for 15 minutes. Total DNA counterstaining was performed using 0.001μg/ml DAPI (4′,6-diamidino-2-phenylindole) for 2 minutes. Afterwards, cells were washed once in 1x PBS solution and a drop of slow fade gold was applied on the cells before covering the cells with a cover glass and sealing with nail polish.

### Microscopy and Imaging

Stained cells were assessed using a Leica TCS SP5 laser-scanning confocal microscope (Leica, Mannheim, Germany). Images were captured with a 63x oil immersion objective. Total DNA (DAPI) was excited with a UV laser at 405 nm, DIC images were captured with an Argon laser (488 nm), while the AlexaFluor 488 conjugated telomere probe was excited by wavelength of 488 nm. All images were captured at a resolution of 1024 × 1024, acquisition speed 200 Hz, and line averaging of 16. Images were sequentially scanned with the aim of generating RGB color images with an 8-bit depth configuration. We only report cells with good overall morphology, passing over cells that were folded, crushed, or otherwise suboptimal and likely representing preparation induced artifacts. Z-stack images were taken with an acquisition speed of 700 Hz (although we initially recorded some Z-stacks with the acquisition speed of 400 Hz), line averaging of 4, and a step size of 0.13 *µ*m respectively. We adjusted the smart gain and smart offset to improve image quality.

### Image analysis

Cell/nuclear volume and total fluorescence intensity were quantified using the ‘Nikon NIS Element’ image analysis software (Nikon, Tokyo, Japan). We manually set points (or outlines) to capture cell size (length x width in *µ*m), and nuclear diameter (by drawing a circle around the nucleus and measuring the diameter in *µ*m) using the measurement tools in ‘Nikon NIS Element’ software to calculate their respective radius (in *µ*m) and volume (*µ*m^3^). Mean intensity of all nuclei was calculated using the ‘ROI (region of interest)’ tool in ‘Nikon NIS Element’ software. Using the ROI tool, we drew a polygonal or circular line surrounding the nucleus, returning the mean intensity using the ‘ROI’ statistics. Total intensity was calculated by multiplying the nuclear volumes (*µ*m^3^) with the mean intensity per pixel.

## Results

We used a laser scanning confocal microscope to characterize nuclear features from a total of 116 individuals stained with both DAPI and a telomere specific probe (Table S1; Table 1). We evaluated cell and nuclear size, volume, total DNA as estimated by DAPI, and amount of gene-sized chromosomes using a telomere probe (Table 1; Fig. 1 and 2). In total, we captured 23 vegetative, 29 conjugating, and 64 developing (early & late developing) cells, which allowed us to infer the life cycle stages (Fig. 3) of *C. uncinata*. Comparing across stages, we found that the conjugating cells tend to be smaller in size, with an average cell size of 35*µ*m, 40*µ*m, and 31*µ*m in length in vegetative, developing, and conjugating stages respectively. (Fig. S9). We also found a number of cells that were not obviously in one of these three categories, and we include them as oddities, which may represent either preparation artifacts or rare events (Figure 1, panel II. a-l).

**Table 1:** Image analysis summary table of *Chilodonella uncinata* cells at different stages during their macronuclear development shows variability in their total DNA and telomere content during their MAC development. Avg= Average; TI= Total Intensity; vol= volume; dev= Developing; MAC= Macronuclei.

| Categorized dev. stages | # of nuclei quantified | Avg. cell vol (µm<sup>3</sup>) | Avg. nuc vol. (µm<sup>3</sup>) | MAC-cell ratio | Avg. TI (DNA) | Avg. TI (Telo) | Avg. DAPI vs. Telo ratio |
| --- | --- | --- | --- | --- | --- | --- | --- |
| Vegetative | 23 | 8171.7 | 168.8 | 0.27 | 24716 | 14122 | 0.63 |
| Early Development | 40 | 7236.48 | 142.4 | 0.20 | 13758 | 6725 | 0.83 |
| Late Development | 24 | 11201.01 | 316.9 | 0.40 | 20566 | 7433 | 0.32 |
| Conjugating | 29 | 5914.46 | 225.4 | 0.48 | 33788 | 22223 | 0.64 |

**Figure 1:**
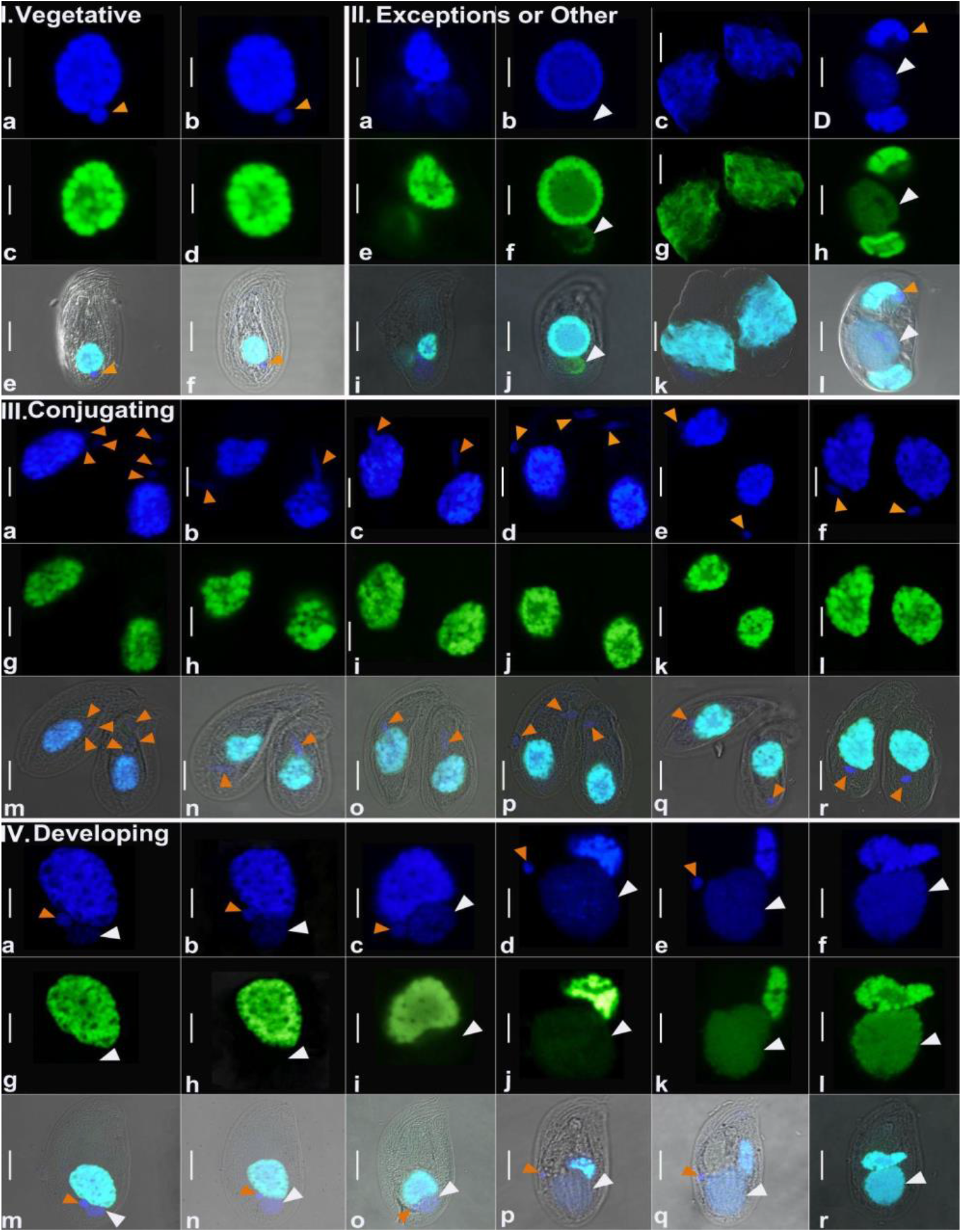
Representative images from each categorized developmental stage, DAPI stained (blue) and Telomere stained (green) nuclei of *Chilodonella uncinata*. **(I)** Representative cells and nuclei from vegetative stage (panel a-f). **(II)** All the exceptional or other forms of nuclei that were captured during the study but not categorized. **(III)** Representative conjugating cells and their nuclei (panel a-r). **(IV)** Representative developing cells and their nuclei; **panel a-c, g-i, and m-o** are representing the early developing stage; **panel d-f, j-l, and p-r** are representing the late developing stage. Orange arrowheads indicate the MICs and white arrowheads indicate the developing MACs. MICs= micronuclei, MACs= macronuclei. Scale bar = 5*µ*m.

**Figure 2:**
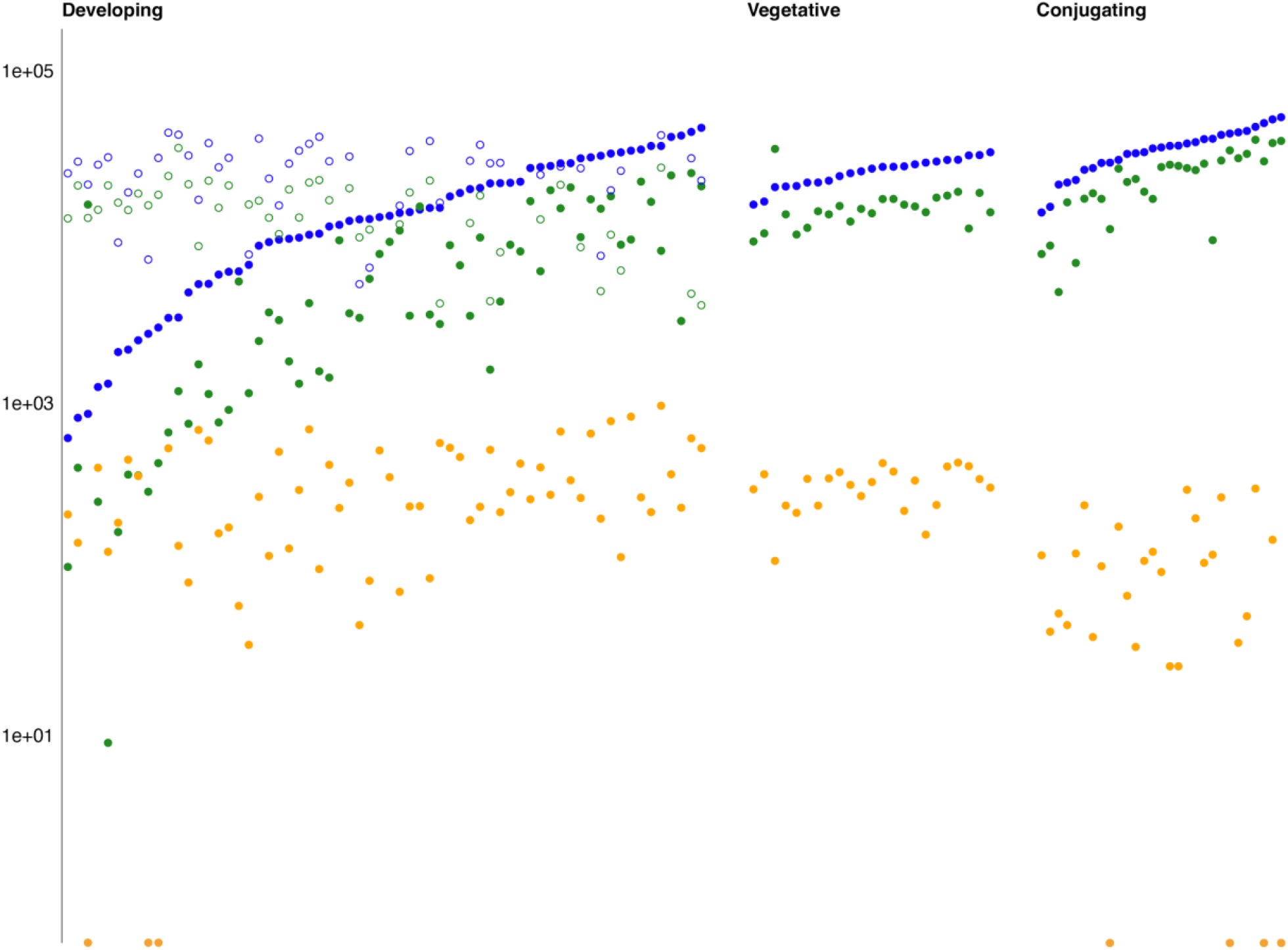
Different stages and total intensity during the macronuclear development in *Chilodonella uncinata* of all categorized cells. X-axis represents the different stages during the macronuclear development in *C. uncinata* and the Y-axis (in log scale) represents their total intensity. Cells are categorized by ‘Developing’ (including early and late developing stages), ‘Vegetative’, and ‘Conjugating’ stages from left to right on the X-axis. Solid blue circles are the newly developing MACs DNA data and solid green circles are the newly developing MACs telomere data respectively. Open blue circles are the old MACs DNA data while open green circles are the old MACs telomere data respectively during early and late development. Solid orange circles indicate the MICs data on the plot. MICs= micronuclei, MACs= macronuclei.

**Figure 3:**
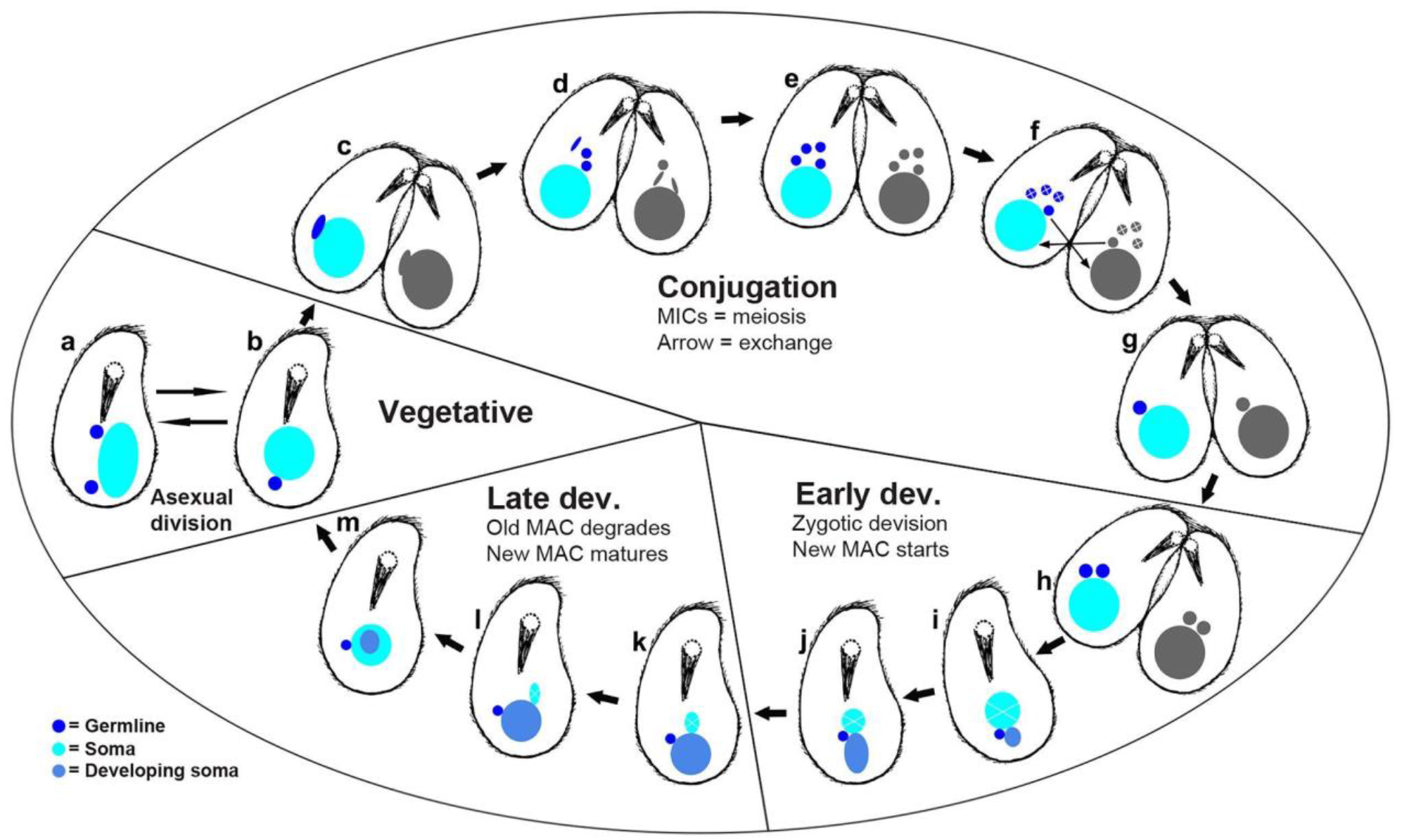
Inferred lifecycle of *Chilodonella uncinata* illustrated by cartoons based on images from Fig 1 and Figs S1-S8.. **(a)** MAC amitosis (asexual division). **(b)** Vegetative cell and nuclei. **(c)** Meiosis-I. **(d-f)** Conjugation, meiosis-II of MICs, and exchange of haploid MICs. **(g-h)** zygotic nuclei and mitotic division of zygotic nuclei. **(i-j)** Early development of MACs. **(k-m)** Late development of MACs. MICs= germline micronuclei, MACs= somatic macronuclei.

### Vegetative cells

We defined 23 vegetative cells analyzed in this study (Fig. S1; Table 1; Table S1) as those containing a large ‘typical’ heteromeric macronucleus (e.g. with densely-stained material surrounding a DNA poor center) with a smaller germline micronucleus below the MAC (Fig. 1 panel I. a-f; Fig. S1 and S2). The vegetative macronuclei stain robustly with both DAPI (blue) and the telomere probe (green) (Fig. 2), consistent with endoreplication of gene-sized chromosomes as seen in *C. uncinata* (Maurer-Alcalá and Katz 2016) *and the congeners Chilodonella cucullulus* and *Chilodonella steini* (Radzikowski 1976, 1985). In contrast, no obvious fluorescence signal for the telomere probe was detected in the germline micronuclei, which are predicted to have only a small number of chromosomes, and hence few telomeres (Fig. 1 panel I. a-f; Fig. S1 and S2).

We observed an almost consistent ratio between total DNA (i.e. DAPI) and telomere signal in the vegetative cells, with only one cell possessing more telomere signal than DAPI (Fig. S1 and S2; Table S1). The consistent ratio between total DNA and telomere coupled with the near doubling of total DNA content as inferred from total DAPI intensity (from 15596 to 32204, Table S1) indicates that these vegetative cells are likely cycling through amitotic division of macronuclei. We did not capture mitosis of micronuclei among these 23 vegetative cells.

### Conjugation

We collected data from a total of 29 conjugating cells, which we identified as pairs of cells joined at their oral apertures (Fig. 1 panel III. a-r; Fig. S3; Table 1). To allow comparisons across life history stages, we choose to measure only one cell, arbitrarily choosing the cell on the right side of all conjugating pairs (Fig. 1 panel III. a-r; Fig. S3). Across all the conjugating cells we analyzed, we were able to capture their meiosis events (both meiosis-I and meiosis-II), plus exchanging of nuclei (Fig. 3c-f; Fig. S4). The ratio between the total DNA (in blue) and telomeres (in green) among all of the cells from the conjugating stage is not consistent, which contrasts with what was observed in the vegetative cells (Fig. 2). In addition, we note that these cells have similar total DNA content as compared to vegetative cells (Fig. 2), despite their smaller size (Fig. S3 and S9).

### Development of somatic macronuclei

Given our interest in nuclear architecture, the bulk of our analyses focused on individuals in various stages of development. After measuring a total of 64 cells determined to be developing based on the presence of both a new and old somatic macronucleus, we inferred developmental stages (Fig. 1 panel IV. a-r; Fig. S5 and S7). We categorized developing cells into two broad subcategories; we define early development as those stages that still contain prominent old MACs, and only weakly-stained new MACs while late development are stages with prominent new MACs. Using these criteria, we ended up categorizing 40 cells in early development and 24 in late development stages (Table 1). We report on DNA and telomere staining of both the ‘old MAC’ (i.e. the one degrading over time) and ‘New MAC’ (i.e. the *anlagen*, or newly-developing nucleus) (Fig. 1 panel IV. a-c; Fig. S5, and S6)

During development, the ratio between the DNA and the telomere content varies, although both increase gradually. By comparing total DNA and amount of gene-size chromosomes between the newly developing macronucleus and the ‘old’ macronucleus (Fig. S10), we observe that the DNA content in the old MACs does not decrease as the new MAC develops (Fig. S10B), though there is a decline in telomere signal (Fig. S10C). This suggests that *C. uncinata* is not recycling old material as it generates a new macronucleus following conjugation (see below).

Based on our detailed observations of thousands of *Chilodonella* cells, with an emphasis on its nuclear architecture across multiple life stages, we propose a revised nuclear life cycle (Fig. 3; Table S1). During conjugation, micronuclei become elongated-shaped and increase in copy number, which we interpret as meiosis I (Fig. 3a). Next, we see additional elongated stages with up to 4 micronuclei per cell, consistent with meiosis II. We inferred that three of these haploid nuclei degrade and the remaining haploid nucleus in each individual are exchanged to each other (Fig. 3.d-f) before undergoing nuclear fusion (karyogamy). The resulting zygotic nucleus undergoes mitotic division; one of the daughter nuclei becomes the new MAC as the other is the new MIC in the mature cell (Fig. 3.g-h).

During *Chilodonella*’s early development, the old MAC retains its ‘typical’ morphology while the newly developing MACs appear attached to them (Fig. 1 panel IV; Fig. 3.i,j, and k-m). We found a consistent positioning of the old and new (developing) MACs during the early and late developing stages: the old MACs tend to be on top (towards the anterior portion of the cell) and the newly developing (new) MACs positions on the bottom (towards posterior portion of the cell-under the old MACs) of the old MACs (Fig. 3.i,j, and k-m; Fig. S5). During the late developmental stage, the old MACs lose their typical morphology and degrade as the new MACs develop with the new MIC attached or almost attached (Fig. 1 panel IV. d-f, j-l, and p-r; Fig. 3.i,j, and k-m). After going through all of these described stages, they return to their vegetative stage (Fig. 3.b; Fig. S1). We recorded a few cases (though we saw many) where the MAC is dividing into two equal parts that we infer as asexual division (Fig. 3.a).

### Exceptions

While scanning thousands of cells under the microscope, we found a few cells with nuclear stages that seemed unusual. One cell contains chromosomes in the newly developing MAC but little or no DNA in the developing MAC (Fig. 1 panel II. b, f, and j), two individuals that had three nodules of MACs with different patterns of its position (Fig. 1 panel II. a, e, i, and d, h,l), and one individual has very densely spread out DNA (Fig. 1 panel II. c, g,k). It is unclear whether these exceptions represent unknown plasticity or dead-ends in the life cycle of *C. uncinata*.

## Discussion

Here we characterize nuclear events in the life-cycle of a non-model ciliate, *Chilodonella uncinata*, using FISH and laser-scanning confocal microscopy. We investigate total DNA with DAPI staining as well as the abundance of gene-sized macronuclear chromosomes (Riley and Katz 2001; Zufall and Katz 2007; Huang and Katz 2014; Gao et al. 2015) with a telomere-specific FISH probe. We report patterns of macronuclear development that differ from those previously described for diverse model lineages, and even for the congeners *C. cucullulus* and *C. steini* (Radzikowski 1976, 1985). We also find unexpected variability in macronuclear DNA content during conjugation, and we present an updated life cycle for this species that extends from a previous study in our lab (Bellec et al. 2014).

Though in some aspects the nuclear processes in *C. uncinata* are similar to model ciliates such as *Tetrahymena, Paramecium, Oxytricha*, we see notable differences in the timing of DNA amplification in the new MAC; we also add data on the relative ratio of total DNA (DAPI) with gene-sized chromosomes (telomere probe). Vegetative stages of *C. uncinata* as expected: DNA content in the heteromeric somatic macronucleus doubles during amitotic division and the ratio of total DNA to gene sized chromosomes remains relatively stable (Fig. 1 panel I. a-f; Fig. S1 and S2; Fig. 2). In other ciliates with extensively fragmented genomes (i.e. possessing gene-sized chromosomes) ratio of total DNA content to gene-sized chromosomes has not been reported by microscopy.

In contrast to the relative consistency in vegetative stages (Fig. 2), the ratio of total DNA to gene-sized chromosomes is much more variable during conjugation, including when meiotic divisions of the germline micronucleus are occurring in *C. uncinata* (Fig. 3c-h). It is unclear why this variability occurs, although it may be in response to the state of the cell and/or the available resources. During development, *Stylonychia* may “recycle” nucleotides from the hyper polyploid somatic nucleus to offset the energetic cost of re-hyperpolyploidizing the newly developing somatic nucleus (Madireddi et al. 1995). However, we do not see the same pattern of recycling in *Chilodonella* as the DNA content of the new MAC increases before the parental MAC degrades (Fig. 2)

The process of MAC development in *C. uncinata* differs markedly from other ciliates with gene sized chromosomes. For example, spirotrich ciliates like *Stylonychia, Oxytricha* and *Euplotes* go through three development stages: (i) an initial amplification stage of the entire genome, (ii) a DNA poor stage (result of massive DNA elimination), and (iii) a final amplification of maturegene-sized chromosomes (Ammermann 1971; Neeb 2016). By contrast, in *C. uncinata*, we observe a concerted increase in the amount of total DNA and gene-sized chromosomes increase during development (Fig. 2). We observe no evidence of clear DNA elimination stage in *C. uncinata*, consistent with the macronuclear development study of this species (Pyne et al. 1974), which suggests that this major step is likely on-going throughout development, resemblant of *Paramecium*’s somatic development (Rzeszutek et al. 2020). Perhaps most surprisingly, we do not see evidence for recycling of nucleic acids, as total DNA content in the old macronucleus does not decline with increasing DNA content in the newly-developing MAC (Fig. 2; Fig. S6, S8); this contrasts with ciliates like *Stylonychia* that appear to reuse nucleotides from the degrading macronucleus to make a new macronucleus (Sapra and Dass 1970; Ammermann 1971; Ammermann et al. 1974; Maercker et al. 1997). Notably, *Chilodonella* is estimated to eliminate only ∼30-35% of its germline, compared to *Stylonychia* that eliminates >90% of its germline genome (Ammermann et al. 1974), which may be a driver for nucleotide recycling in the latter species.

We present an updated life cycle of *C. uncinata* that incorporates the data we collected by fluorescence microscopy (Fig. 3). Here we show changes in the position of the meiotic products prior to conjugation, and of both the newly-developing MAC and MIC during development. We observed changes in the MIC position during developmental events, with the MIC repositioning from the posterior region of the cell to being nestled between the old and developing somatic nuclei. Most studies of nuclear developmental in other ciliates report that the developing MAC is often positioned towards the posterior portion of the cell, which we also report in *C. uncinata* (Zhang et al. 2023; Ishida et al. 1999; Jurand et al. 1964; Sapra and Dass 1970; Neeb 2016; Gong et al. 2020). Together these data from *Chilodonella uncinata* expand our knowledge of ciliate life cycles.

## Acknowledgements

We are grateful to Judith Wopereis (Center for Microscopy and Imaging - Clark Science Center, Smith College) for help with the confocal microscopy and image analysis process. We used open AI in generating several R scripts used in this study. This work was supported by grants R15HG010409 from the NIH and OCE-1924570 from NSF to L.A.K.

## Author contributions

Ragib Ahsan-designed research, performed research, analyzed data, and wrote the paper; Xyrus X. Maurer-Alcalá-writing: review & editing; Laura A. Katz-designed research, performed research, analyzed data, writing: review & editing

